# Ensemble Refinement of mismodeled cryo-EM RNA Structures Using All-Atom Simulations

**DOI:** 10.1101/2024.07.24.604258

**Authors:** Elisa Posani, Pavel Janoš, Daniel Haack, Navtej Toor, Massimiliano Bonomi, Alessandra Magistrato, Giovanni Bussi

**Affiliations:** Scuola Internazionale Superiore di Studi Avanzati (SISSA), Via Bonomea 265, 34136 Trieste, Italy; National Research Council of Italy, Institute of Material Foundry at International School for Advanced Studies (SISSA/ISAS), via Bonomea 265, 34136 Trieste, Italy; Department of Chemistry and Biochemistry, University of California, San Diego, La Jolla, CA, USA; Institut Pasteur, Université Paris Cité, CNRS UMR 3528, Computational Structural Biology Unit, Paris, France

## Abstract

The advent of single-particle cryogenic electron microscopy (cryo-EM) has enabled near-atomic resolution imaging of large macromolecules, enhancing functional insights. However, current cryo-EM refinement tools condense all single-particle images into a single structure, which can misrepresent highly flexible molecules like RNAs. Here, we combine molecular dynamics simulations with cryo-EM density maps to better account for the structural dynamics of a complex and biologically relevant RNA macromolecule. Namely, using metainference, a Bayesian method, we reconstruct an ensemble of structures of the group II intron ribozyme, which better match experimental data, and we reveal inaccuracies of single-structure approaches in modeling flexible regions. An analysis of all RNA-containing structures deposited in the PDB reveal that this issue affects most cryo-EM structures in the 2.5—4 Å range. Thus, RNA structures determined by cryo-EM require careful handling, and our method may be broadly applicable to other RNA systems.

---

RNA molecules play a vital role in cells by acting as both carriers of genetic information and catalysts (ribozymes) to regulate gene expression, proteome diversification and, ultimately, to perform protein synthesis [1–3]. RNA function is often dictated by its 3D structure, and this is particularly true when considering catalytic RNA molecules or proteins/RNA and small-molecules/RNA interactions [4]. With the advent of single-particle cryogenic electron microscopy (cryo-EM) techniques, it is now possible to resolve large macromolecular complexes, including RNA-containing systems, with a resolution comparable with that of traditional X-ray diffraction experiments [5–9]. However, the number of solved RNA structures remains limited in comparison with protein structures. The standard approach for the analysis of cryo-EM data consists in classifying millions of acquired 2D single-particle images into structurally homogeneous 2D class averages, and then reconstructing one or more 3D density maps. The following step, namely generating a 3D model that fits the obtained density map, usually assumes that a single 3D structure can reproduce the density associated with a very large number of 2D images. For high-resolution maps (*<* 3 Å), structures can be built by using the same software employed for X-ray crystallography [10, 11]. However, for cryo-EM maps with resolutions *>* 4 Å, *de novo* model building is also adopted [12–16].

All the methods mentioned, as single structure approaches, aim at reproducing a cryo-EM potential density map by building a single model. However, applying these approaches to heterogeneous and dynamic biomolecules, where the observed density maps might result from a mixture of structures (conformational states), is highly challenging. This is particularly true for disordered or multi-domain proteins and for RNA systems [17, 18]. Fitting such a potential density map with a single structure might indeed lead to non-biologically-relevant models or structural artefacts if the employed density map originates from a mixture of heterogeneous conformations. In these approaches, structures are normally refined by performing energy minimizations. Accurate MD simulations can also be used for cryo-EM structure refinement [19, 20]. Standard approaches are based on empirical potentials enforcing the atoms positions of the simulated structure to match the measured density maps [21–27]. Although accounting for some conformational variability in the space confined to the EM map potential, the MD simulation is here used to search for a global minimum. Importantly, the conformational landscape may become very rugged and feature multiple proximal local minima, possibly causing the fitted structure to become “trapped” within local minima and resulting in structurally poor or functionally irrelevant models. This problem can be tackled using a replica-exchange approach [28]. More advanced methods allow an automatic inclusion of conformational averaging [29, 30]. Among these methods, metainference-based MD simulations [31] can be fruitfully used to reconstruct structural ensembles. Specifically, metainference allows to enforce the agreement between the experimental density map and an average computed simulating a number of replicas of the system. The replica averaging procedure results in a conformational ensemble that is as close as possible to the underlying MD ensembles [32, 33] and, at the same time, minimizes model discrepancy with respect to experiment [31]. Additionally, the dynamical nature of the refinement process allows to include soft constraints that can enforce independently available structural information. The application of cryo-EM-guided metainference to refine structural ensembles of macromolecular complexes has so far been limited to proteins [30, 34–38]. Importantly, this method has never been tested on large RNA molecules.

Here we apply cryo-EM guided metainference-based MD simulations to the group II intron ribozyme from *Thermosynechococcus elongatus*, a ∼ 800-nucleotide long RNA macromolecule that can function as a catalytic RNA by performing self-splicing reactions and by acting as a retroelement through its insertion into double-stranded DNA [39]. Besides its biological significance as evolutionary ancestor of the spliceosome and its potential application as a gene editing tool, this structure was selected because it is one of the few cryo-EM RNA-mostly structures deposited in the PDB data bank, and was obtained using a single-structure refinement procedure [40]. We reveal that, for such a highly plastic system, a single structure cannot be at the same time compatible with the experimental data and with the expected structure of base-paired RNA helices, resulting in mis-modeling of some regions. For this system, the remodeling mostly affects flexible regions, which are located in the exterior, solvent exposed stem loops and are not phylogenetically conserved. In contrast, functional domains involved in catalysis are well-ordered, exhibit high-quality cryo-EM density, and are phylogenetically conserved. As a result, these core catalytic regions of the ribozyme are relatively rigid and did not require remodelling. Interestingly, a detailed analysis of all the deposited RNA-containing structures revealed that the modeling problem with flexible helical regions broadly applies to RNA-containing cryo-EM derived structures in the 2.5–4 Å resolution range.

## Results

### Test system and preparation

First, we visually inspected the deposited structure of the group II intron ribozyme (PDB code: 6ME0, cryo-EM map: EMD_9105, resolution 3.6 Å) [40]. The secondary and tertiary structures are reported in Fig. 1 A and C. A 38-nucleotide long gap was present, which we modeled using DeepFoldRNA [42] (see Methods). In addition, a number of supposedly canonical RNA helices could be visually seen to be not properly paired. A subsequent more systematic analysis, based on a combination of annotation [43] and secondary structure prediction [44] (see Methods), confirmed the presence of six helices that should be present based on the predicted secondary structure, but are not properly folded in the solved structure (Figure 1 and Table S1). We then run 2.5 ns-long molecular dynamics (MD) simulations in explicit solvent (see Methods) to remodel correctly these helices by restraining them to the the structure of canonical template RNA duplexes with the same sequence (Figure 1B). To this aim we used the ERMSD metric [41], which accounts for the presence of properly paired strands and has been used to reversibly fold RNA motifs [45] or to model structures starting from a coarse-grained prediction [46].

**Figure 1:**
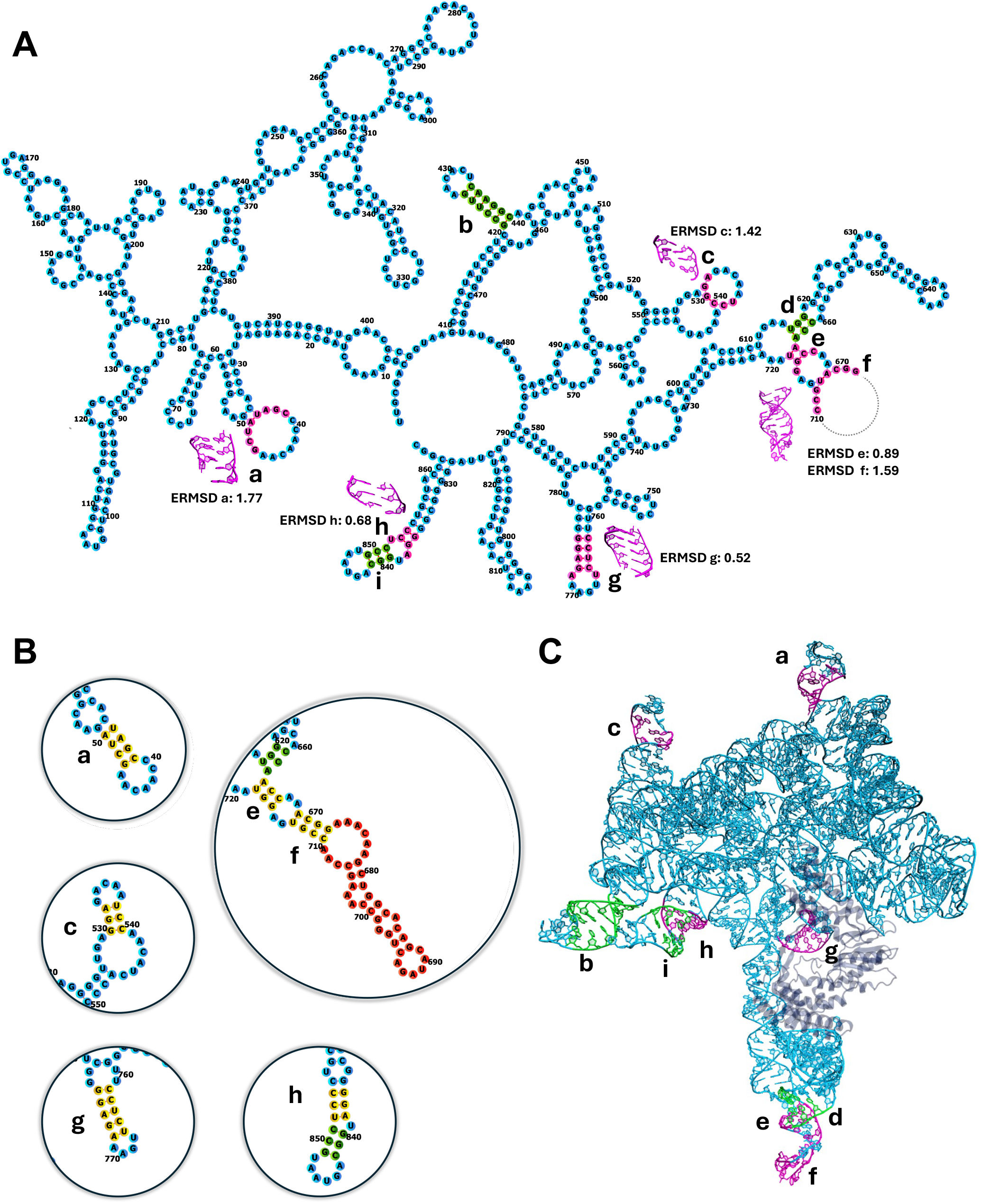
**A**, Secondary structure of the group II intron from *Thermosynechococcus elongatus*, as obtained by annotating the deposited PDB structure (6ME0). Helices object of investigation are labelled as (a-i). Regions predicted as base paired by secondary structure prediction tools, but that are not properly paired in the PDB are highlighted in magenta. The 3D structure of these regions as reported in the deposited PDB is also shown in magenta. Deviations of the deposited helices from an ideal helix are measured using the ERMSD metric [41] and reported. Base pairs that were correctly formed in the PDB structure, but that unfolded during the all atom simulations are shown in green. The dashed line indicates the missing part in the PDB structure, close to helix f. **B**, The regions properly paired after remodeling are enclosed in circles, with the corresponding base pairs highlighted in yellow. The missing domain, modeled with DeepFoldRNA [42], is shown in red. **C**, 3D structure as deposited in the PDB structure (PDB code 6ME0) with the helices highlighted using the same colors and numbering of panel A. The protein part is depicted in transparent blue.

### Ensemble refinement

After having rebuilt a complete structure with properly folded helices, starting from the final structure of the restrained MD simulation, we performed cryo-EM guided metainference simulations [31], to maximise the accordance with the experimental density map while accounting for the ribozyme plasticity. According to a well-established protocol devised for proteins [36], we initially performed a single-replica refinement in which we enforced one single model to match the experimental data. As expected, when this single-replica refinement was run without any restraint on the six remodeled helices, these quickly unfolded. This confirms that the properly folded helices are not directly compatible with the experimental density within the single-structure assumption. Conversely, when restraining their helicity, whitin this single-replica refinement approach, we observed the unfolding of three additional helices (helices b, d, and i in Figure 1) possibly due to other structural constraints. The only way to preserve the folding of all helices listed in Figure 1 during the single-structure refinement was to explicitly restrain all of them. Next, we performed ensemble refinement metainference simulations by using an increasing number of replicas (8, 16, 32, and 64) and collecting a 10 ns long trajectory for each replica. Interestingly, a minimum number of 8 replicas was required to satisfy the experimental restraints in the metainference MD simulation. Specifically, all the simulations attempted with a lower number of replicas were crashing reporting missing convergence in enforcing bond constraints, which indicates that the experimental and helical restraints were mutually incompatible. This observation *per se* is suggestive of the substantial dynamics of this ribozyme. Helical restraints were released after the first 5 ns, and only the remaining part of the MD simulations trajectories was used for the analysis.

The most representative structures extracted from the 32-replicas simulations (Figure 2B) exemplifies the inherent plasticity of this macromolecular complex. The most flexible region is the *ab initio* modelled loop, in agreement with the fact that its structure was eluding experimental determination in the single structure refinement procedure. However, also other parts of the ribozyme display significant dynamics, as was expected from the high values of B-factors, shown in Figure 2A. The flexibility of the different regions of the ribozyme can be also visualized in Figure S9, where the root-mean-square fluctuations is reported.

**Figure 2:**
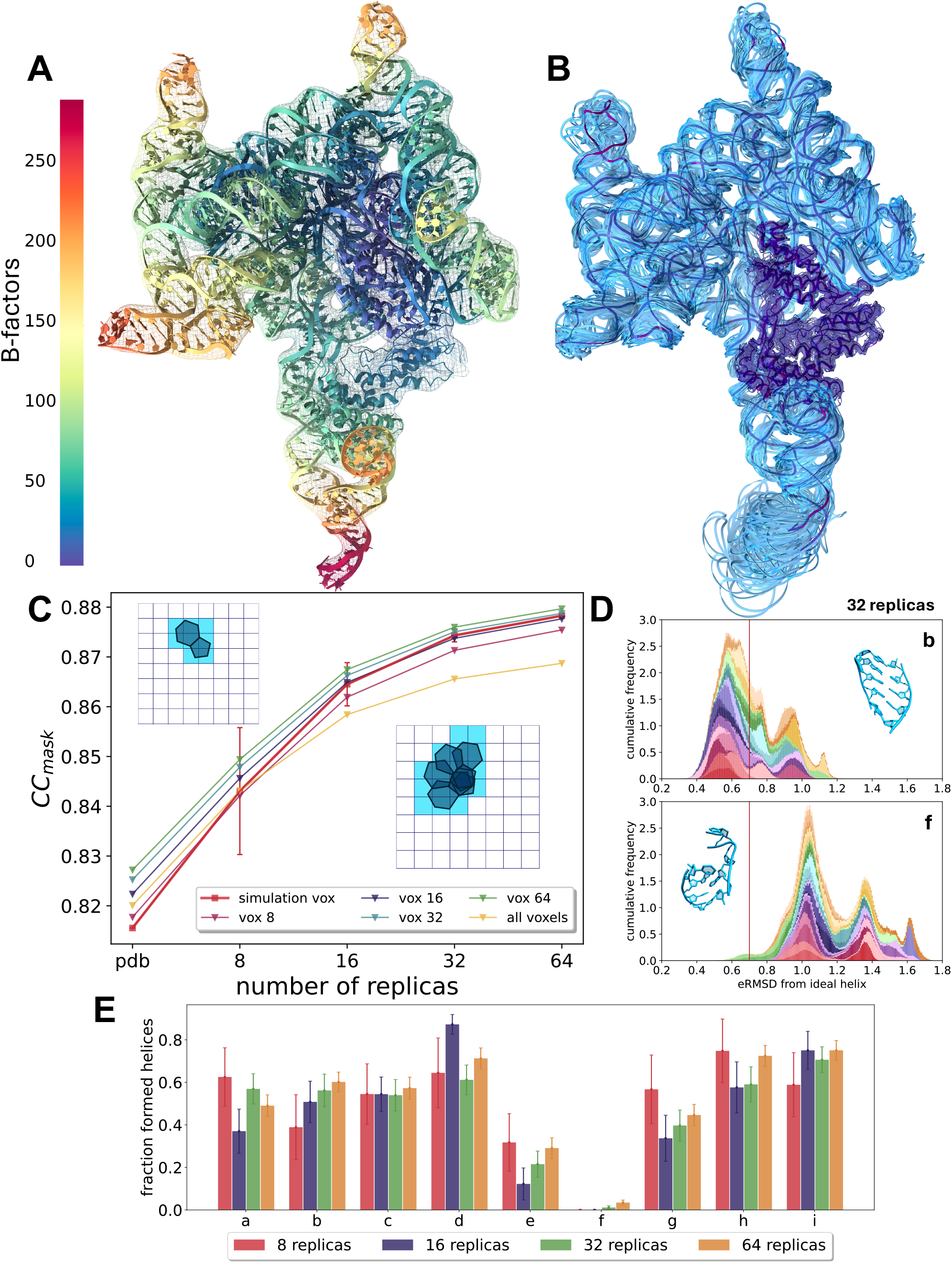
**A**, PDB structure colored by B-factor values (increasing value from blue to red) to highlight the most flexible regions of the structure. **B**, centroids of the 32 replica simulations (blue), together with the original PDB structure (purple), that display dynamics in the higher B-factor external helices and in the modelled gap part. **C**, Accordance with the experimental potential density map, tested with different sets of voxels. The voxels were selected around the initial structure (simulation vox) using all the voxels explored during the replica simulations with 8, 16, 32 and 64 replicas, or including all the voxels. The accordance was monitored for the PDB structure and the multiple-replicas simulations (8, 16, 32 and 64). **D**, Examples of the ERMSD distributions in the 32-replica simulation for helix b, which displays both folded and unfolded conformations, and helix f, which is always unfolded. The red line indicates the standard threshold for ERMSD value to discern the two states [43] **E**, Fraction of ensembles from the replica simulations with formed helices during the second part of the simulation, where helix restraints were removed.

### Back-calculation of density maps

The trajectories obtained with metainference simulations were then analyzed by back-calculating the corresponding averaged density map and comparing it to the experimental one (Figure 2C). The cross-correlation coefficient (CC_mask_), which accounts for agreement between the calculated and the experimental map inside the mask calculated around the macromolecule [47], clearly increases with the number of replicas, irrespective of the exact definition of the region used to do the comparison. This is expected since the agreement with the experiment is enforced and the flexibility of the model increases with the number of replicas. The standard error of the CC_mask_ between the predicted and experimental density markedly decreased with an increasing number of replicas, consistently with the corresponding increase in statistics. Indeed, as discussed below, most of the conformational heterogeneity originates from the differences between replicas. As a result, 32 replicas appeared to be the best compromise between agreement with experiment and computational cost.

### Analysis of helical regions

Next, we monitored the folding of the nine helices that were restrained in the initial part of the metainference simulation. The CC_mask_ computed only on those helical regions was significantly larger than the global CC_mask_ (Figure S1), suggesting that, when properly modeling helical flexibility, the experimental density map could be correctly reproduced.

The actual structure of the nine enforced helices was quantitatively analyzed by computing their ERMSD [41] with respect to their corresponding template A-form helices. An analysis of the cumulative distribution of the ERMSD value sampled in the 32-replica-simulation (Figures S2–S5) revealed that some of them retained a highly dynamic behaviour even in the metainference MD simulation. Sample distributions are shown in the main text for helix b, which adopted both a folded and unfolded state (Figure 2D) during the simulation, and for helix f, which never folded. Figure 2E displays the fraction of structures with a properly folded helix, for every helix and in each of the simulation setups. Most helices retained the ideal folded state in more than 50% of the ensemble, with helices e and f showing the least percentage of folded structures. However, these helices are placed nearby the gap in the structure that we modelled *ab initio* (Figure 1C). Therefore they lie in the region of the map with low signal, which corresponds to a high dynamics of the structure.

### Convergence of the simulations

In order to assess how much the percentage of ideally folded helices was depending on the length of the trajectory we also prolonged the 32 replica metainference simulations to 20 ns per replica, achieving very similar results (Figure S6). This finding might be surprising, considering that the time scale of the metainference simulation (10 ns per replica) was much shorter than both the time scale actually required to observe these fluctuations in RNA molecules [17] and the typical time scale spanned by MD simulations [48]. This choice was mostly dictated by the high computational cost associated with the on-the-fly back-calculation of the cryo-EM map needed for enforcing agreement with experiment and the necessity to run a large number of replicas in parallel. To verify that a 10 ns long MD simulation within the metainference scheme was indeed sufficient to sample the structural dynamics of the investigated system, we made a systematic comparison of the generated structures, both intra replica and inter replica. Specifically, we performed a full annotation [43] of the structures produced during (a) a reference 1 *µs* MD simulation and (b) during the metainference simulation, and quantitatively estimated their heterogeneity (see Methods). Results for the 32-replica simulation and for the plain MD simulation are shown in Figure 3, other simulations are analyzed in Figure S7. The structural pair-wise distance between pairs of snapshots reports on the difference in their pairing and stacking pattern. As a result, the typical distance between structures produced in different replicas is significantly larger than the typical distance between any pair of structures produced in the long MD simulation. In fact, the typical distance between structures produced in the 1 *µs* long MD simulation is comparable to the distance observed between structures produced in the same replica of the 10 ns long metainference MD simulation. Therefore the structures extracted from different replicas appear to be more heterogeneous than those sampled from the much longer plain MD simulation. This analysis shows that the use of replicas has a major benefit on sampling and confirms the effectiveness of the multi-replica metainference approach in sampling the structural ensembles of flexible biomacromolecules. To further support this finding, we evaluated the CC_mask_ of the 32 replicas using a single representative frame for every trajectory (i.e. the centroid of each MD simulation replica, Figure 2B). As a result, we obtained a CC_mask_ value of 0.87, which is compatible with the value obtained analyzing the whole trajectories (see Figure S8). We also tested different lengths of the simulations, obtaining the same result for 1, 5 and 15 ns. Conversely, the CC_mask_ value of the 1 *µs* unbiased MD simulation (see Methods), was only 0.33.

**Figure 3:**
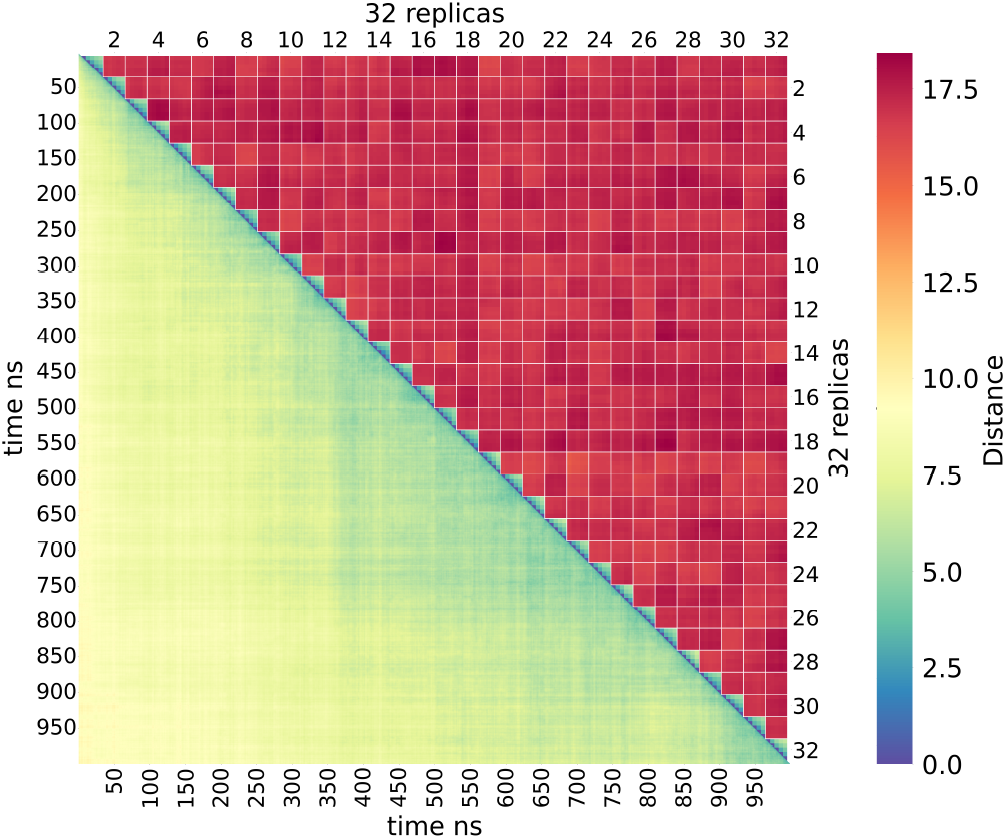
Annotation distances among the 32-replica MD simulations (each 5 ns-long). Intra- and inter-replica distances are reported along the diagonal and in the upper triangle, respectively. Distances of parts of a 1 *µs* long MD simulation used as a reference are shown in lower triangle.

### PDB-wide analysis of incorrectly folded RNA helices

To assess the general impact of the canonical single-structure assumption in refining RNA structures, we performed a systematic analysis of all RNA-containing structures determined by cryo-EM currently deposited in the Protein Data Bank (PDB). Specifically, we inspected all the cryo-EM structures with resolution *<* 6 Å that contained more than 100 nucleotides-long RNA strands. Structures were extracted and annotated as discussed in Methods. Some of the structures present in the PDB database were discarded due to issues in the annotation or incomplete nomenclature in the PDB database. Hence, this analysis covers 1296 structures (full list in the associated GitHub repository).

For each structure, we performed a secondary structure analysis equivalent to that performed on the structure 6ME0. Namely, a secondary structure prediction was done using as constraints the annotated base pairs, and the number of nucleotides that were predicted to be paired, but that were not paired in the annotation, was calculated, and then normalized over the sequence length. Figure 4A shows a scatter plot comparing the reported resolution of each of the analyzed structures with the fraction of nucleotides in which the secondary structure prediction and the actual annotation of base pairing were not corresponding. The result is expected to be dependent on the specific annotation tool employed, and might falsely report unpaired nucleotides in cases where the secondary structure prediction tools fail in identifying the most stable structure. However, we observed that even structures in the 2.5–4 Å high resolution range have a very significant fraction of nucleotides that are annotated as unpaired. Examples of improperly folded helices belonging to these high resolution structures are shown in Figure 4B, where it can be seen that Watson-Crick hydrogen bonds are not correctly formed.

**Figure 4:**
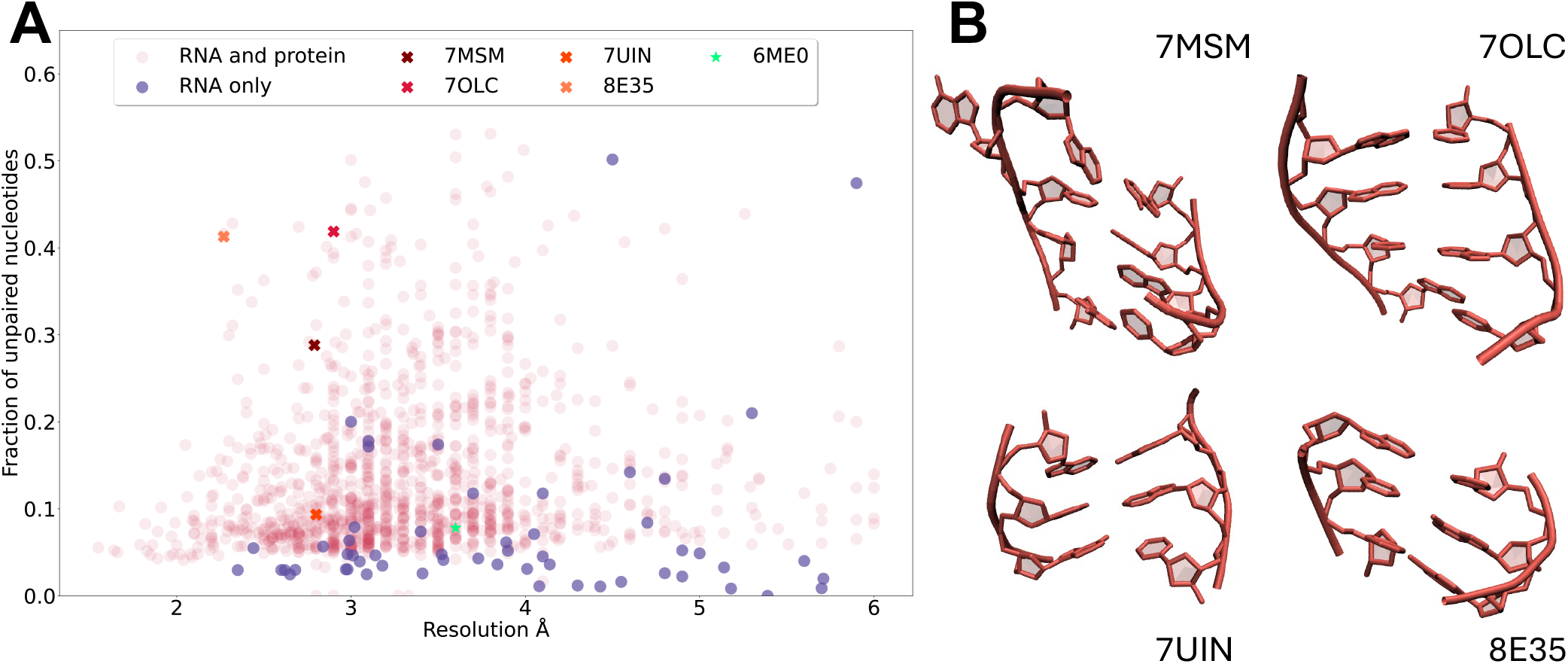
**A**, Structural analysis performed on 1296 deposited cryo-EM structures from the Protein Data Bank, with resolution *<* 6 Å. The structures have been analysed to predict base pairings that are not present in the deposited PDB, but that could exist based on the sequence. The total number of unpaired nucleotides was then divided by the length of the sequence (the values range from 0 when all the base pairings are formed, to 1 if no base pairs exists). The blue data points represent RNA-only structures, while the pink ones are RNA-protein complexes. The group II intron ribozyme investigated here (6ME0) is highlighted in green and examples of PDBs where many unformed base pairs are found (7MSM, 7OLC, 7UIN, 8E35) are marked with Xs. **B**, Examples of mis-modeled base parings, in more pathological cases (7MSM) and less serious ones (8E35), taken from the aforementioned PDBs.

## Discussion

In recent years, remarkable advances in single-particle cryo-EM have enabled the structure determination of an increasing number of bio-molecular assemblies with unprecedented detail and resolution. Nevertheless, although cryo-EM maps can frequently reach the atomic-level resolution, the information that they provide often covers a wide range of resolutions across the entire density map, with conformationally variable regions having a substantially decreased resolution. These low resolution regions often correspond to flexible domains. In these cases, the ensemble of coexisting conformations contributing to the measured cryo-EM images poses challenges to structural biologists. Indeed, the common approach of single structure fitting to cryo-EM potential density maps may result in artifacts due to flexible domains.

Here, we have evaluated the difficulties of using cryo-EM maps of inherently dynamic RNA macromolecules for structural determination. As a prototypical example we chose the group II intron ribozyme, considering an experimental structure that was previously derived using single-structure refinement [40]. This specific structure is representative as it covers a range of relevant cases. The core of the molecule is very well structured and solved at high resolution, while a highly flexible peripheral region could not be determined from the experimental map. At the same time, several helices were in regions considered rigid enough to be accurately determined by single-structure refinement but, after careful analysis, resulted to be incorrectly paired in the deposited PDB structure. After having modeled the missing region and enforced the correct pairing in the helical regions, we proceeded with the systematic application of cryo-EM based metainference simulations with an increasing number of replicas (8, 16, 32 and 64), which showed better agreement with the experimental data. Our results clearly indicate how the structural heterogeneity underlying the plasticity of this large ribozyme can not be accounted for by the single PDB structure, which we showed to have a lower CC_mask_ with experiment when compared to the generated ensemble. Indeed, an ensemble refinement based on 32 replica simulation represents for this system a good trade-off between agreement with the experimental data, correct helix modeling, and computational cost of the simulation. The 32 representative structures are also available for download in the associated GitHub repository. We notice that the idea of combining molecular dynamics simulations and experiments to reconstruct the structure of dynamical RNAs is not new. However, this has been so far done on much smaller RNA molecules, mostly using nuclear-magnetic-resonance or scattering data (see [49] for a recent review). Cryo-EM data has been traditionally combined with MD within the single-structure approximation.

In spite of its relatively large computational cost, mostly related to the need to simulate the system in explicit solvent and to back-calculate the cryo-EM map on-the-fly during such simulation, the computational approach used here is ultimately cheap when compared with state-of-the-art unbiased MD simulations, which typically covers the multi-microsecond timescale, yet being less efficient in accounting for the conformational variability of this RNA macromolecule. This is due to the conformational heterogeneity guaranteed by the presence of multiple replicas and of the restraint on their average, which allows to obtain converged results in relatively short simulation timescales. Hence, even relatively short simulations can be used to generate heterogeneous ensembles provided that enough replicas are simulated, making the approach highly parallel and suitable for high-performance computing setups. Importantly, all the simulations were performed using open source software and all the relevant input scripts and files are made available, so as to facilitate the application of our protocol to other RNA systems.

We suggest the resulting ensemble is a more faithful representation of the dynamics of the group II intron compared to the deposited single-structure model. Interestingly, not all the helices that we initially enforced were formed in the final ensemble, suggesting that some of them might have been incorrectly predicted by the thermodynamic model used here or that their population could be too low to be observable. Overall, the combination of helical restraints and cryo-EM based metain ference simulations provides an ensemble that optimally combines physical modeling, previously known information, and experimental data.

It is worth mentioning that the group II intron deposited in [40] had two dinstinct structures (6ME0 and 6MEC). These structures correspond to separate conformational states from the same cryo-EM dataset. Hence, the analysis performed in Ref. [40] was already able to correctly split the two conformations. The remodeling performed here is complementary to this process and works at a finer scale: once the major conformations have been classified, the local dynamics pertaining to each of them can be resolved with molecular dynamics simulations.

Most importantly, a systematic analysis of the protein data bank, revealed that most of the RNA-containing or RNA-only structures deposited therein are affected by modeling artefacts. Indeed, we observed many long mismodeled helices at an intermediate resolution range. Namely, 45% of the RNA-containing structures present mismodeled helices larger than 5 base pairs, and 8% larger than 10. This is most likely due to fitting single-structure models into cryo-EM density maps collected from highly dynamic biomolecules. Furthermore, the use of simplified force fields might also contribute to mismodeling artefacts. These structures must be therefore handled with care, and the computational approach presented here offers an effective and viable way to significantly improve them. Importantly, the precise results of this analysis might depend on the annotation method used. Here we used Barnaba [43], which has been optimized to analyze X-ray structures and might be slightly too restrictive in identifying correctly folded helices. However, the visual inspection of randomly chosen structures clearly shows that modeling RNA helices constrained to cryo-EM maps is affected by the single-structure assumption. We notice that Auto-DRRAFTER [16], a recently developed tool for modeling RNA molecules based on the observed cryo-EM density map, is designed to take into account the secondary structure of the modeled system and, in this sense, should be able to achieve a good performance in modeling helical regions. One could therefore envision using these models as starting point for ensemble refinement with our proposed approach. Unfortunately, we were not able to test Auto-DRRAFTER on our system because the current version is limited to modelling RNA-only systems.

We also remark that, given the growing number of cryo-EM structures in the structural databases and the fact that these models are routinely used to train artificial-intelligence methods to predict new structures from sequence, artefacts in the deposited models could easily propagate to new predictions. We tested the recently released AlphaFold3 [50] on the group II intron ribozyme, but unfortunately the predicted structure was completely different from the deposited one, making the comparison very difficult. We stress that the performance of AlphaFold3 on systems where RNA-RNA interactions are predominant is currently under investigation [51]. Additional explorations in this direction, for example using cryo-EM data to refine initial models generated by artificial intelligence approaches, are certainly a new avenue that we believe should be further explored in the future. Furthermore, we envision the possibility to use reweighting methods based on the direct analysis of cryo EM images to further refine the ensembles generated with our approach [52, 53].

In summary, we have illustrated the application of metainference, an ensemble refinement technique based on all-atom MD simulations, to a large RNA-containing macromolecular complex. The accurate physico-chemical models of RNA and the surrounding environment result in a higher computational cost with respect to the fast model prediction of previously reported and more recent deep-learning based algorithms. However, our approach can provide an accurate structural characterization of RNA systems at atomic level, while accounting for its dynamics, a key requirement for broadening our understanding of RNA functions. Furthermore, in the future one could envision generating several structural ensembles of RNA molecules and complexes using our approach and then train a deep-learning model to predict ensembles from cryo-EM maps in a more computational efficient way, thus enabling large-scale determination of accurate RNA ensembles.

## Data availability

All the input files and centroids corresponding to the 32 replicas are available at on GitHub https://github.com/ElisPo/Cryo-EM-refinement/. Metainference input files are available on the PLUMED NEST (plumID:24.016) [54]. Full trajectories for the metainference simulations, excluding solvent, are available on Zenodo at https://zenodo.org/doi/10.5281/zenodo.12761120.

## Online methods

### Model Building

We built our model of the group II intron ribozyme starting from the model deposited in the PDB (id: 6ME0). Since this structure presents a gap of 38 nucleotides, we modelled the missing part with the online webserver of DeepFoldRNA [42], using default settings and the fasta sequence of the missing part (residues 673 to 710) with 17 additional nucleotides for a total of 55 nucleotides (complete sequence from 663 to 717: ACCAAACGGAAACAAGCUGGCACAGCAUAGACUGGGCCAAAGCCAACCGUGAGGU). A portion larger than the missing gap was modeled in order to facilitate the alignment and attachment of the modelled part to the rest of the structure. We also estimated the secondary structure of this sequence with RNAfold and checked that the output of DeepFoldRNA was consistent with the RNAfold prediction [44]. We then aligned the nucleotides in common with the PDB structure and merged the two structures. We also added the 2 ^*′*^ -5 ^*′*^ lariat bond between residues U1 and A860, using Amber tleap [55]. Finally, we ran a minimization *in vacuo* with GROMACS-2021.5 [56].

### System setup and Molecular dynamics (MD) simulations

To model the ribozyme at physiological conditions we replaced the Na^+^ ion in the PDB structure with a K^+^ ion. We inserted the model in a rhombic dodecahedron periodic box and solvated with 398121 water molecules, 1563 K^+^ ions and 788 Cl^*-*^ ions to reach charge neutrality at physiological salt concentration of 150 mM. We then run a new minimization with the solvent. The system topology was built with the OL3 force field (FF) for RNA [57], OL15 FF for DNA [58], both with the bsc0 corrections [59], and ff14SB FF for the protein [60]. Joung-Cheatham parameters were used for the monovalent ions [61], while Li-Merz parameters were used for the Mg^2+^ [62]. Water molecules were described using the TIP3P model [63]. All the simulations were performed using GROMACS-2021.5, at constant temperature T=300 K [64]. Plain MD simulations were performed at constant pressure p=1 bar [65], whereas metainference simulations were performed at constant volume. We used LINCS bond constraints [66] and treated electrostatics with particle-mesh Ewald [67].

### Identification of new base pairings

In order to identify possible base pairings that are not present in the deposited PDB, we started from the original structure 6ME0. We annotated the Watson-Crick base pairings present in the original structure using Barnaba [43] in dot-bracket notation. Then, we run a prediction of possible additional base-pairings not present in the original structure with RNAfold [44]. Here we enforced in the prediction all the base-pairings already found ((*·*)). Since this tool does not handle pseudo-knots, we prevented any bond with all the bases involved in this kind of bonds from the annotation ([*·*],{*·*}, *< >*). From this prediction, we were able to find new base pairings that were missing in the original structure, and correspond to the helices shown in Figure 1 and Table S1.

### Helix restraints

After having identified the mismodeled helices lacking the predicted base-pairings, we generated ideal helix models by using fd_helix (https://casegroup.rutgers.edu/fd_helix.c). Starting from the energy-minimized model described in the previous section, we performed an MD simulation with restraints on all the mismodeled helices (Figure 1). To this end we used the ERMSD [41] metric as implemented in PLUMED [68], which accounts for both the distance between nucleobases and the orientation, and we forced the ERMSD value of the simulated helix to assume a value of 0, corresponding to that of the ideal helix of the same sequence, with harmonic constant 500 kJ/mol. We ran a 2.5 ns long restrained simulation to attain a ERMSD < 0.7 for all targeted helices. The final value of ERMSD from the target helices was < 0.3. These simulations were performed with no additional restraint and in the constant pressure ensemble. Two structures are supposedly equivalent when the ERMSD value is below the threshold of 0.7 [43].

### Plain molecular dynamics

Starting from the structure with all the targeted helices in their ideal conformation, we performed a reference 1 *µs* long MD simulation, at constant pressure and without any restraint.

### Metainference simulations

To perform the metainference simulations, we followed the procedure described in [36] and at https://github.com/COSBlab/EMMIVox. This method requires a modified version of PLUMED 2.9 and will be available in the coming version 2.10. Namely, we started from the structure with all the targeted helices in their ideal conformation and we selected the experimental density map within 3.5 Å from the starting model. We decided not to use the entire map since the computational cost of the simulations would have been much larger without a concrete improvement of the results. Using this cropped map, we initially ran a single-structure refinement for 10 ns, enforcing one single conformation to match the experimental data. In this single-structure refinement step, we sampled the B-factors using a Monte Carlo approach to maximize the accordance with the experimental data. During this step, we also kept the restraints on the helices. The single-structure refinement resulted in the unfolding of 3 additional helices (Figure 1, in green). Therefore we performed again the refinement protocol by adding restraints also to these 3 helices. The refined model that retained all properly folded helices was the starting point of the metainference multi-replica simulations. Multiple metainference simulations were then performed using a different number of replicas: 8, 16, 32 and 64. The initial frames of each replica were selected from the single-structure refinement trajectory as equally spaced. In the first 5 ns of each simulation, we kept the restraints on all the helices in Figure 1 to prevent them from unfolding. In the second part, we removed the restraints to allow the system to explore the conformational space without additional restraints beside the cryo-EM density map. In all the cryo-EM guided simulations, agreement with density maps was enforced every 4 steps to decrease the computational cost [69].

### Accordance with the experimental data

In order to assess the accordance of the simulated structures with the experimental data, we followed the protocol described in [36]. We generated a map for each frame of the trajectory and evaluated the average map along the trajectory. We then performed a linear regression among the voxels of this map and the corresponding ones in the experimental potential density map and we calculated the Pearson correlation coefficient (here called CC_mask_). In order to assess the robustness of the results, we used different sets of voxels for the calculation of the CC_mask_. Namely, we selected (i) the voxels around the initial structure, which are those where we enforced the experimental data during the simulations; (ii) all the voxels explored during the simulations of the replicas (8, 16, 32 and 64, separately); (iii) all the voxels in the map. To test a single structure, we estimated the map that would be generated from the structure, limited to the selected voxels.

### RMSF

During the single structure refinement, we estimated new B-factors using a Monte Carlo approach, needed to maximize the accordance with the experimental density map. Then, for each ensemble-refinement simulation, we compared the average RMSF per nucleotide, among the metainference replicas, with the RMSF obtained from the B-factors determined during single-structure refinement using:

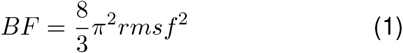

From Figure S9, it can be seen that the RMSF values do not depend on the number of replicas used during the simulation. In addition, there is a high correlation between the RMSF values obtained from the single-structure refinement B-factors and those calculated from the metainference ensembles. As shown visually in Figure S9, the targeted helices display peaks in the RMSF, denoting a greater dynamics. This is particularly true for the modelled part in the gap. Interestingly, this confirms the hypothesis that the corrupted density in that region is due to a higher mobility of the molecule.

### Distances

To assess the distances between the replicas, we started by dividing the second part of the trajectory of each replica (after removing the restraints) into 5 parts, each 1 ns long, and annotated each frame using Barnaba. For every part of the trajectory, we counted how many times each base pairing [70] and stacking (using the Barnaba definition) occurred, thus obtaining a histogram of interactions along the trajectory. We evaluated the distances between parts of the trajectories by calculating the Euclidean distance between the corresponding histograms divided by number of frames.The same procedure was applied to the unbiased MD simulation trajectory, that we instead divided in 1000 parts of 1 ns each.

### Base pairing analysis of the Protein Data Bank

For the systematic analysis of the base pairs in the experimental structures deposited in the PDB database, we developed an automatized protocol based on the following steps: (i) download of the structures (PDB or cif format), later converted into PDB format with AmberTools ccptraj [55]; (ii) annotation of the Watson-Crick base pairings present in the structure using Barnaba in dot-bracket notation, (iii) writing the sequences of the RNA molecules in FASTA format; (iv) prediction of possible additional base-pairings not present in the original structure with RNAfold. As explained previously, we enforced in the prediction all the base-pairings found and prevented any bond with all the bases involved in pseudoknots. In addition, if there were gaps in the structure, we added a UUUU sequence to make the structure continuous and prevented RNAfold to pair these nucleotides. From the prediction, we were able to propose new base pairings that were missing in the original structure. Since some structures presented a very low number of base pairings, and RNAfold thus did not have many restraints and was free to do a prediction that could be not precise, we computed the distances of the new predicted base pairings in the original experimental structure. As a result, it appears that ∼80% of predicted base pairs are in a position in accordance with a pairing (N1/N9 distance < 15 Å), as shown in Figure S10. Most probably this analysis slightly over-estimates the problematic modelling of helices, but it can still give a perception of how problematic the helix modeling in cryo-EM derived RNA structures can be.

## Acknowledgements

Samuel Hoff (Institut Pasteur) is acknowledged for useful discussions. This work has been funded by the Next Generation EU project PRIN 2022 (2022Z4FZE9), EMBO Scientific Exchange Grant (10033) and EURO-HPC (2023R03-136). A.M. and P.J. thank PNRR: National Center for Gene Therapy and Drugs based on RNA Technology CUPB83C22002860006 CN0000004.

