## Supplementary Material for "Ensemble Refinement of mismodeled cryo-EM RNA Structures Using All-Atom Simulations"

| helix name | residues | PDB ERMSD |
|---|---|---|
| a | 36 37 38 39<br>50 49 48 47 | 1.77 |
| **b** | 422 423 424 425 426 427<br>439 438 437 436 435 434 | 0.41 |
| c | 530 531 532<br>540 539 538 | 1.42 |
| **d** | 616 617 618<br>663 662 661 | 0.32 |
| e | 664 665 666<br>718 717 716 | 0.89 |
| f | 669 670 671 672<br>713 712 711 710 | 1.59 |
| g | 762 763 764 765 766<br>775 774 773 772 771 | 0.52 |
| h | 836 837 838<br>854 853 852 | 0.68 |
| **i** | 840 841 842<br>851 850 849 | 0.41 |

Table S1: Identified misfolded helices from the annotation of the deposited PDB structure 6ME0. Residues involved in the creation of the helix and ERMSDs evaluated from comparison with ideal helices with the same sequences of the examined ones, before the refinement. The additional helices that unfold during the simulation (see following paragraph) are in boldface.

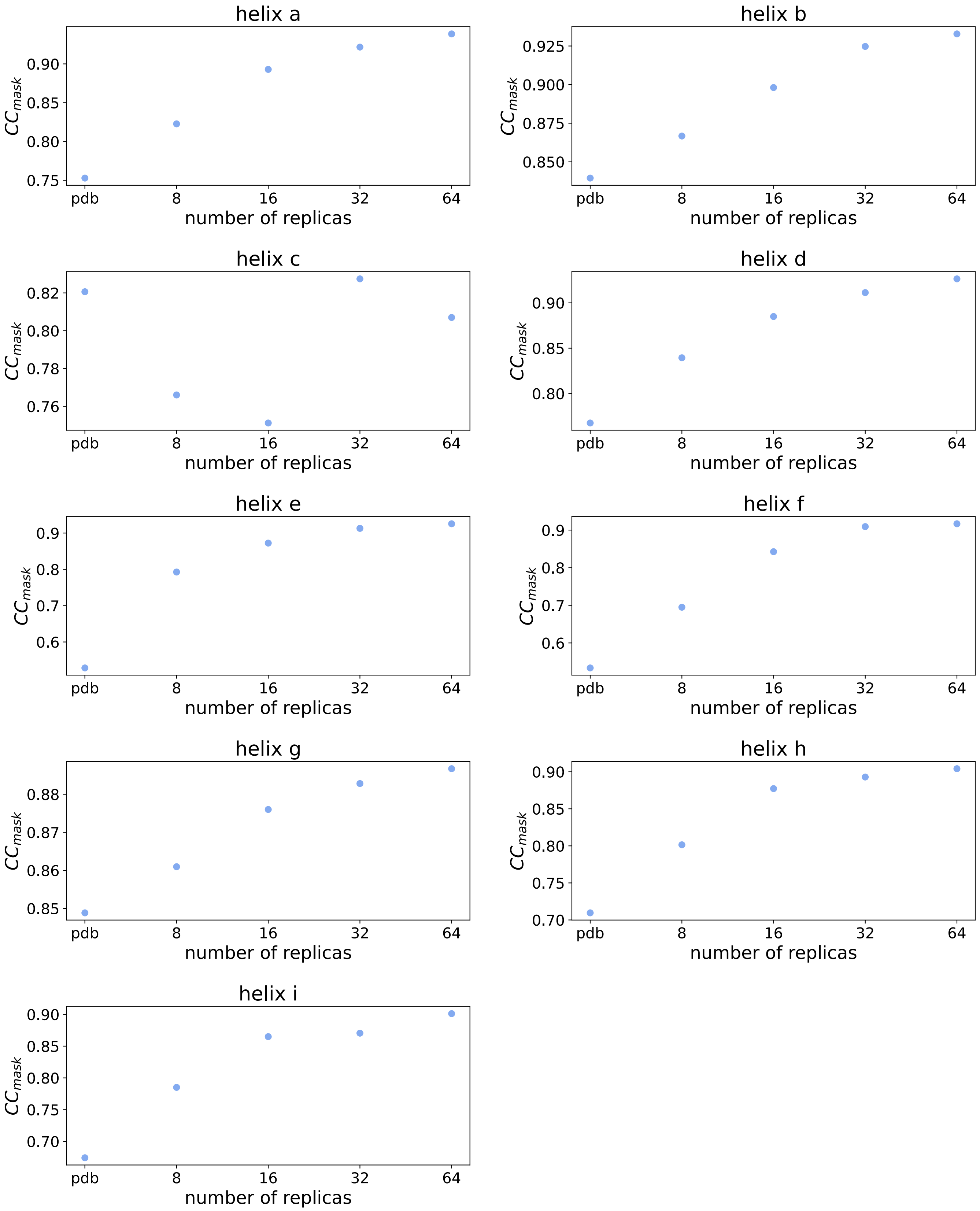


Figure S1: $CC_{mask}$ values for the different setups (deposited PDB, 8, 16, 32 and 64 replicas) evaluated only on the voxels surrounding the 9 targeted helices, separately.

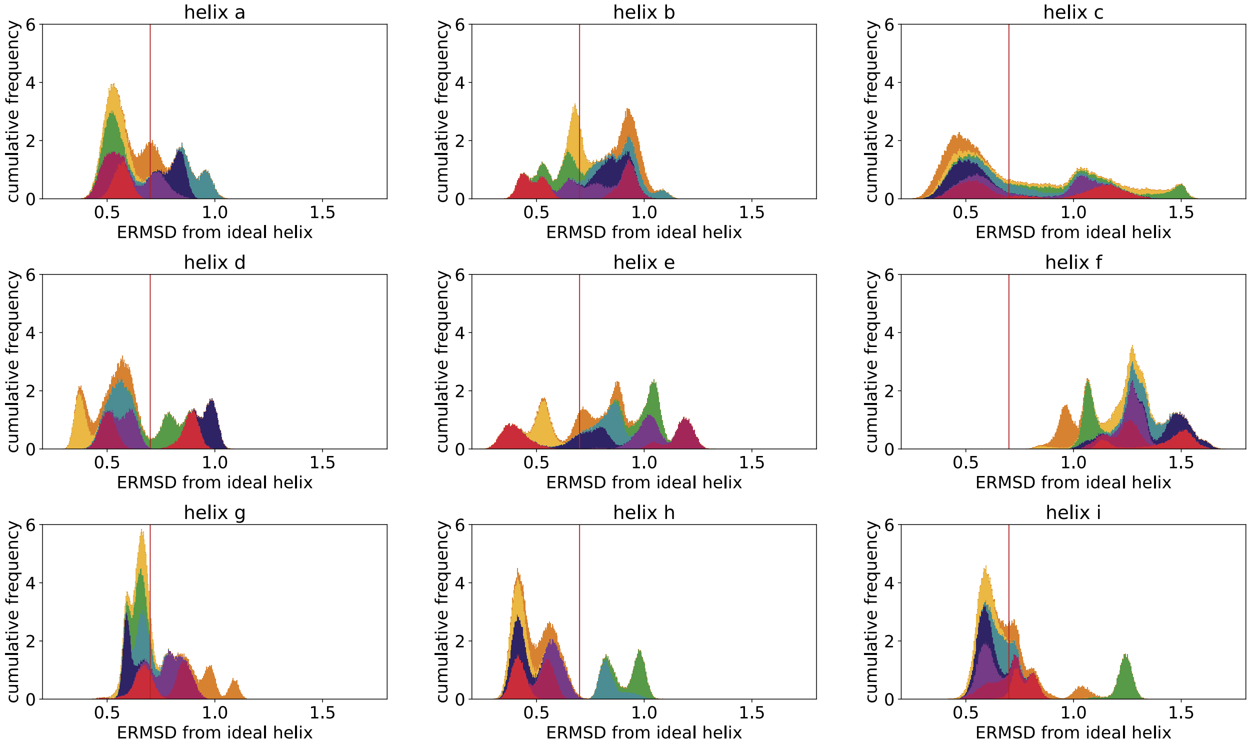


Figure S2: Cumulative distributions of ERMSDs for the 9 restrained helices in the 8 replica simulation.

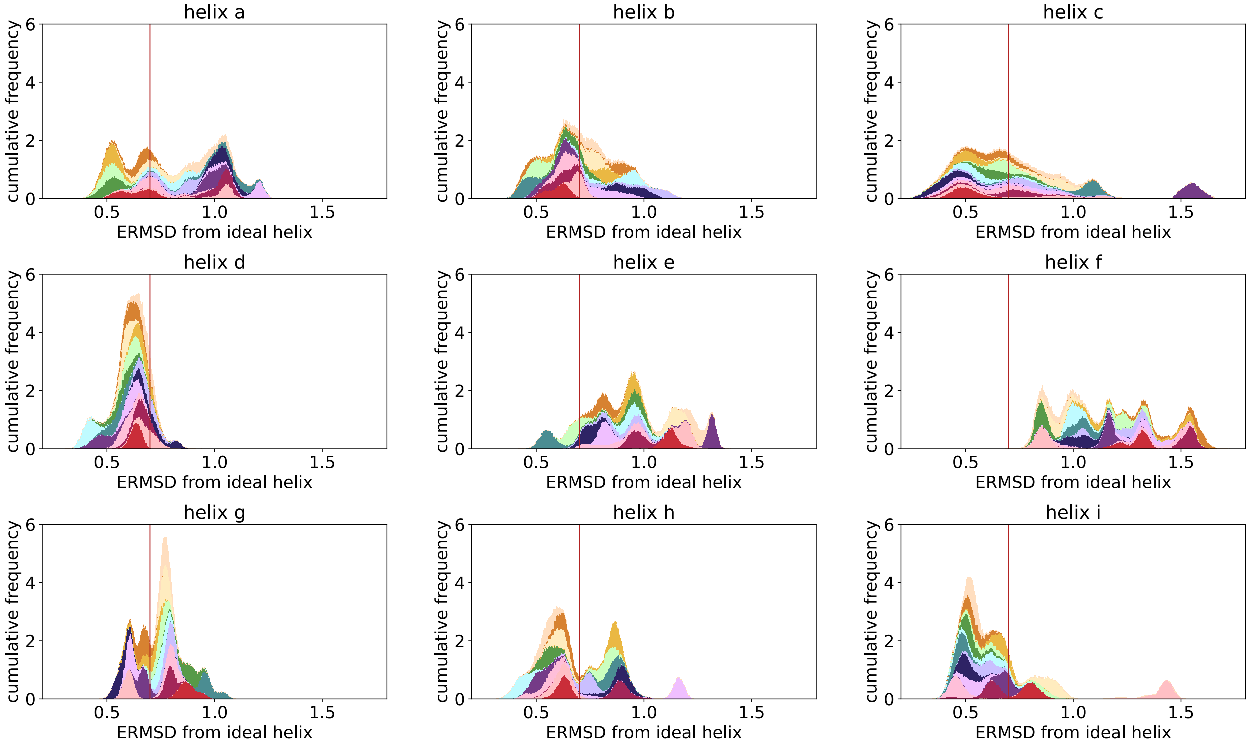


Figure S3: Cumulative distributions of ERMSDs for the 9 restrained helices in the 16 replica simulation.

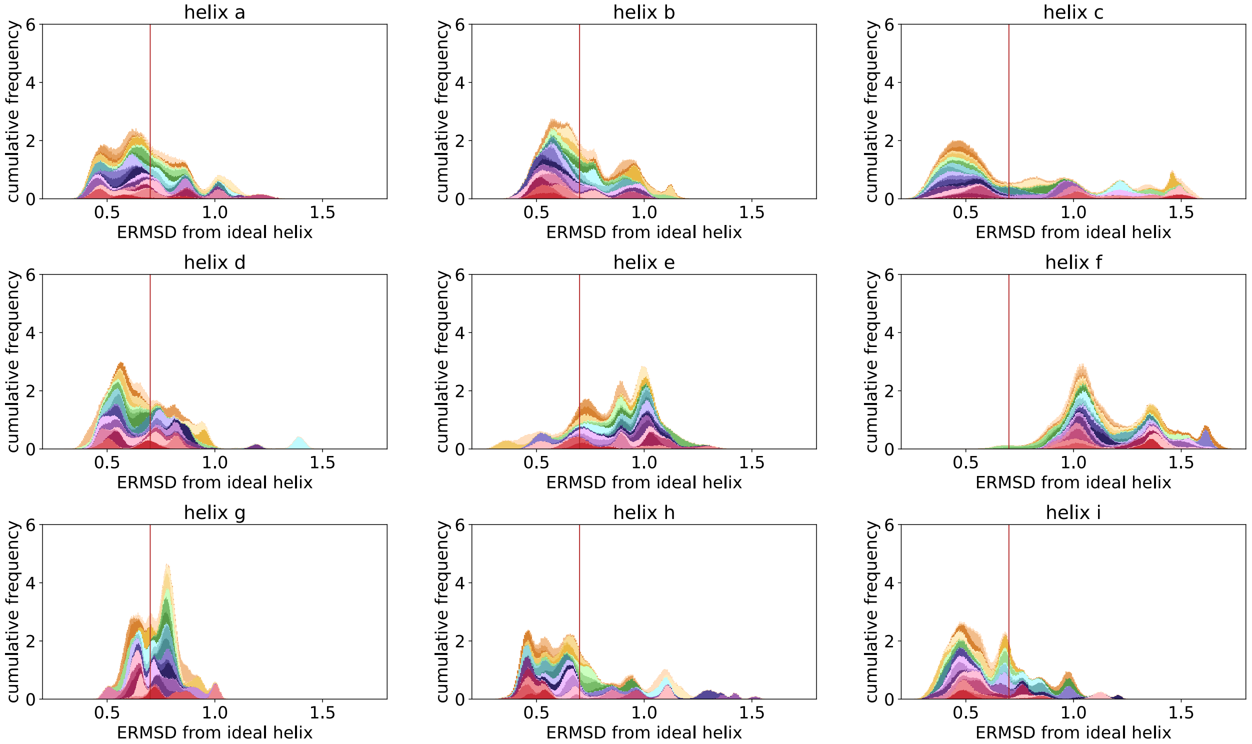


Figure S4: Cumulative distributions of ERMSDs for the 9 restrained helices in the 32 replica simulation.

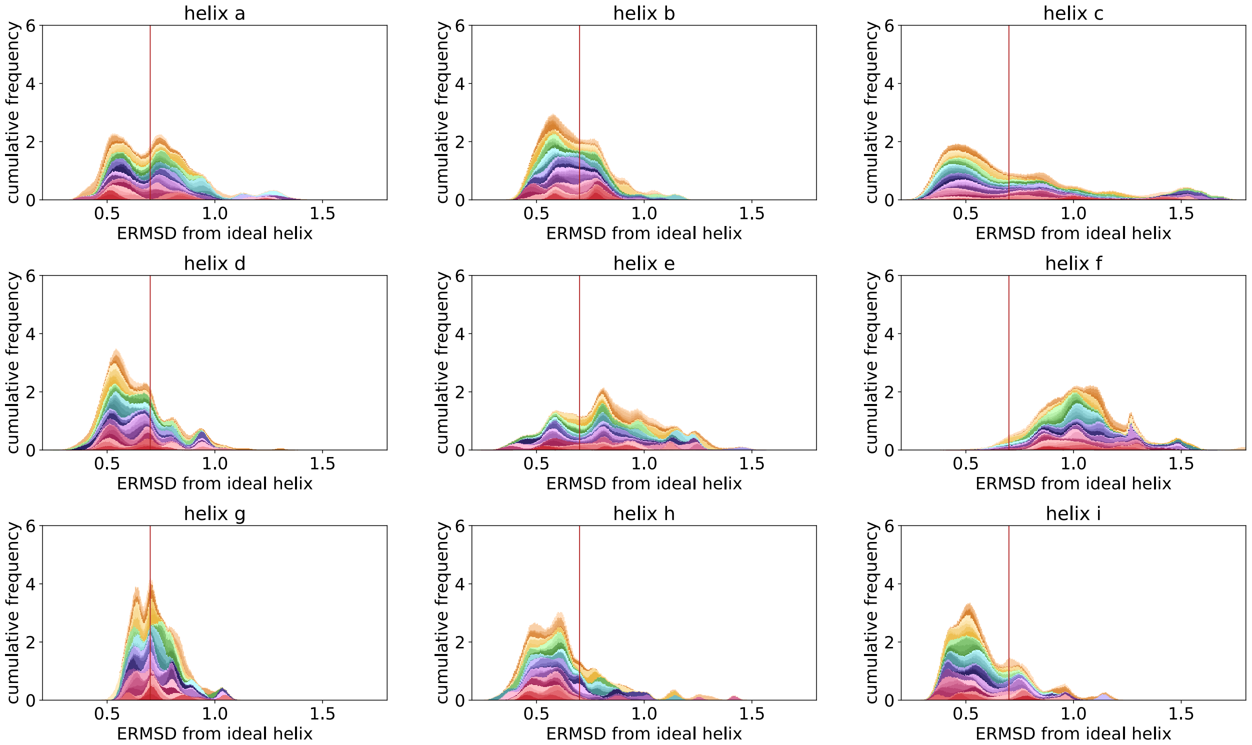


Figure S5: Cumulative distributions of ERMSDs for the 9 restrained helices in the 64 replica simulation.

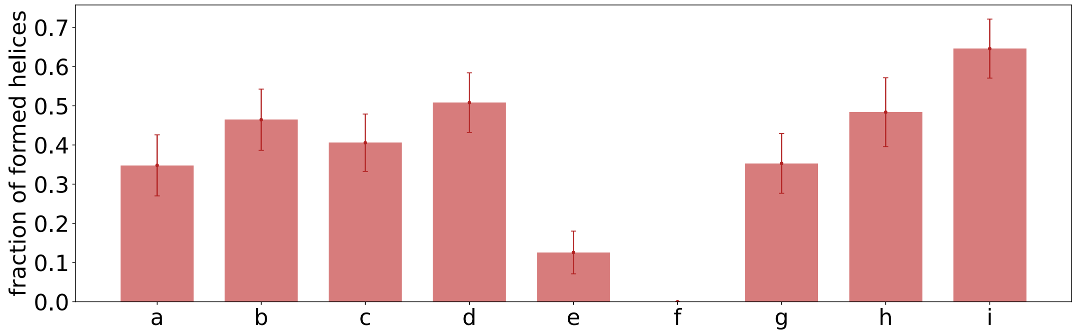


Figure S6: Fraction of folded helices in a 15 ns simulation, after 5 ns of simulation keeping the restraints on the helices, only for the 32 replica setup.

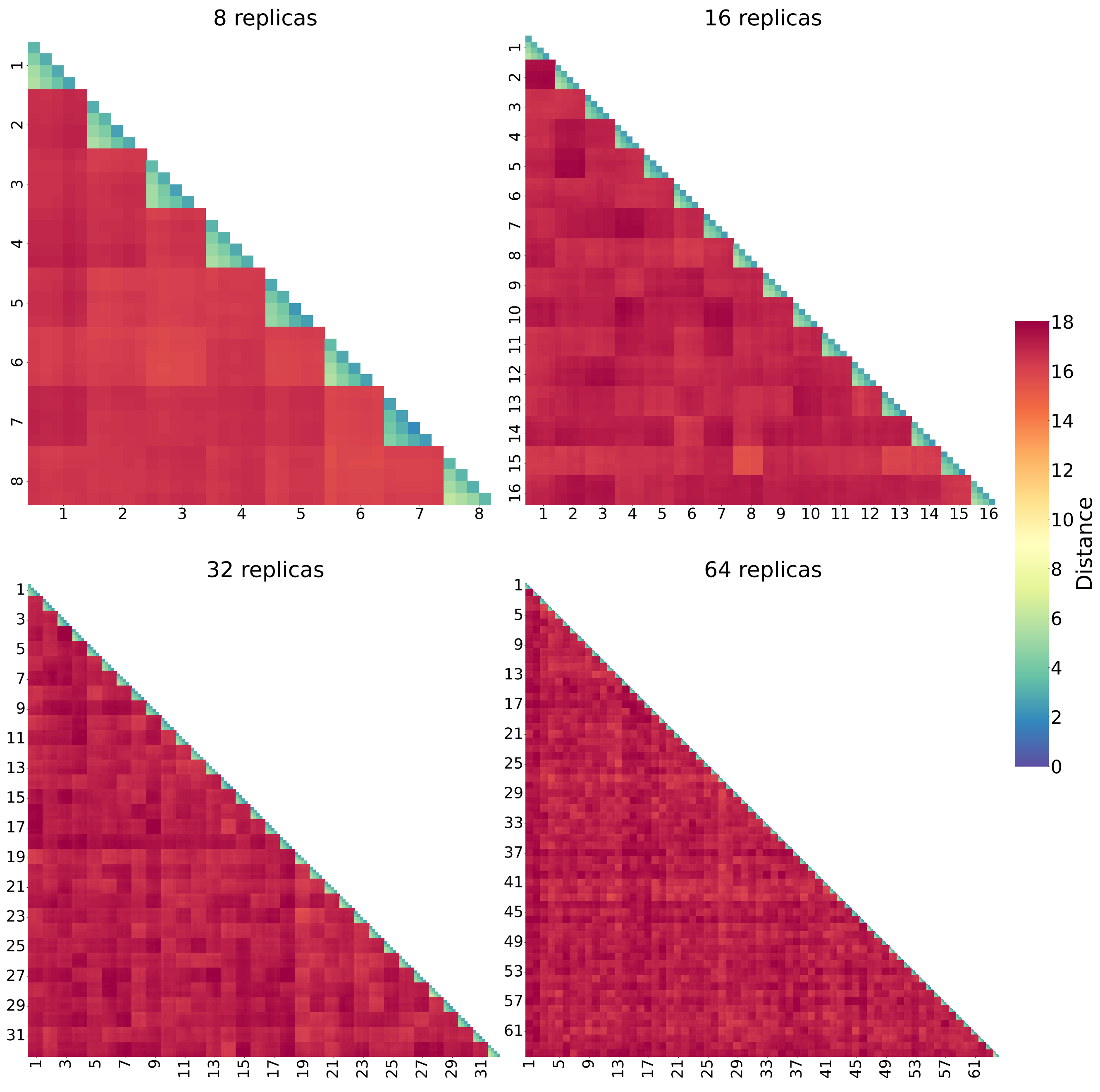


Figure S7: Distances among 5 parts of each replica trajectory, for all the replica setups (8, 16, 32 and 64 replicas). Each part is 1 ns long. Along the diagonal there are the intra-replica distances, while the other squares are inter-replica distances.

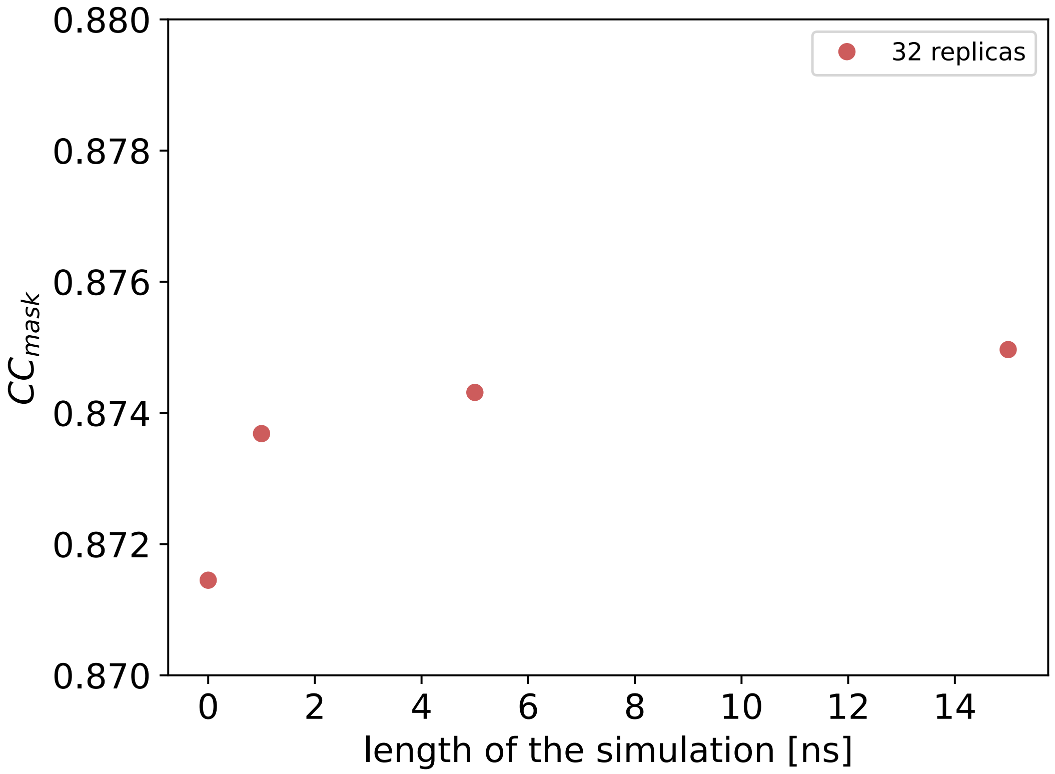


Figure S8: Values of $CC_{mask}$ evaluated on the 32 replicas, using different length of simulations: 0 accounts for only the centroids, the first 1,5 and 15 ns of simulation. The value increases in the different tests, but not significantly.

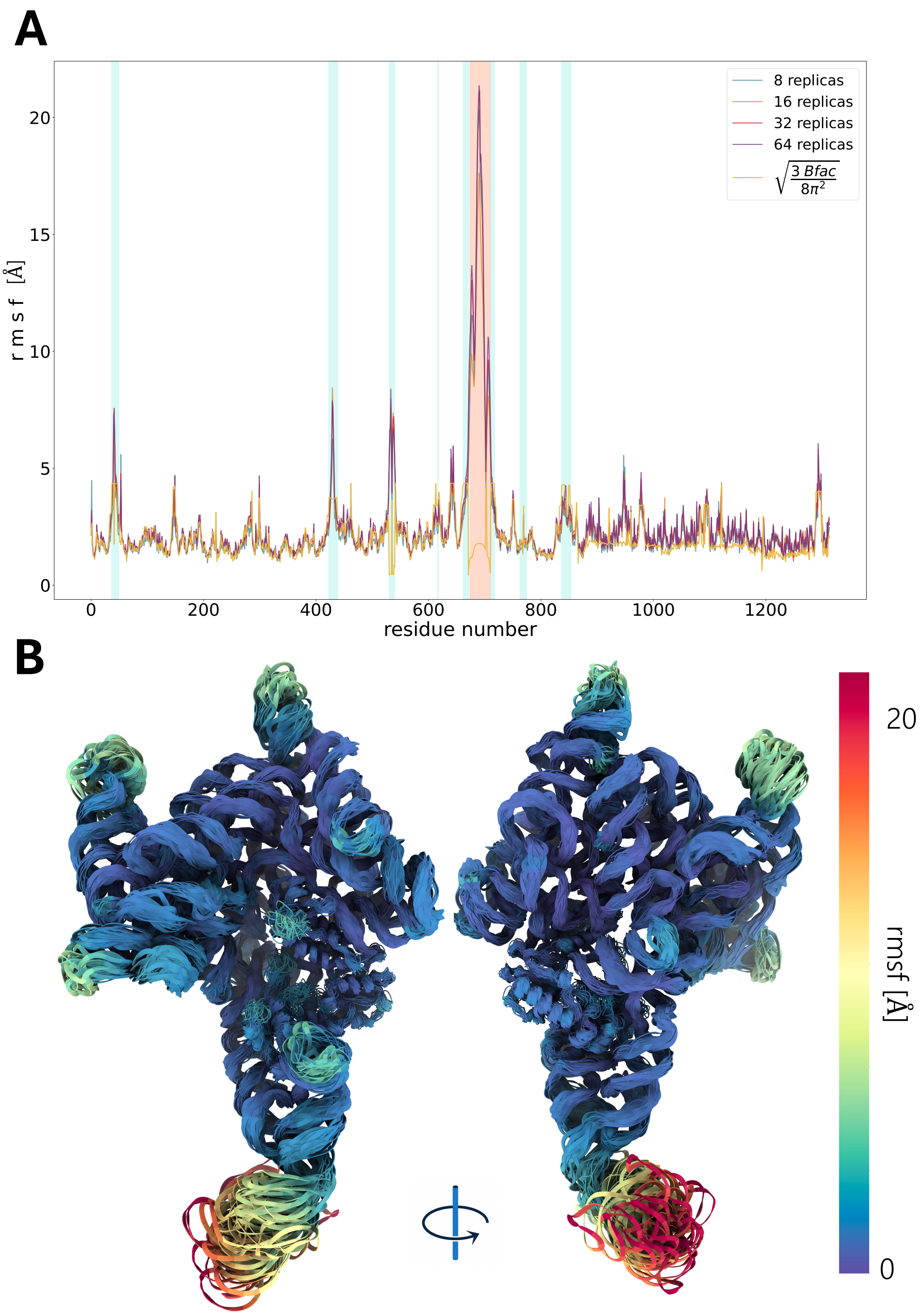


Figure S9: **A**, Average RMSF per nucleotide of the structure during the metainference simulations with 8, 16, 32 and 64 replicas. The value does not depend on the number of replicas. Also the values of the estimated B-factors (see Methods) are shown, transformed into RMSF-scale with equation 1, and appear highly correlated. The targeted helices are highlighted with a turquoise background, while the gap region in orange. **B**, Centroids of the 32-replica simulations, with residues coloured by average rmsf, as reported in A. The two sides of the structures are shown.

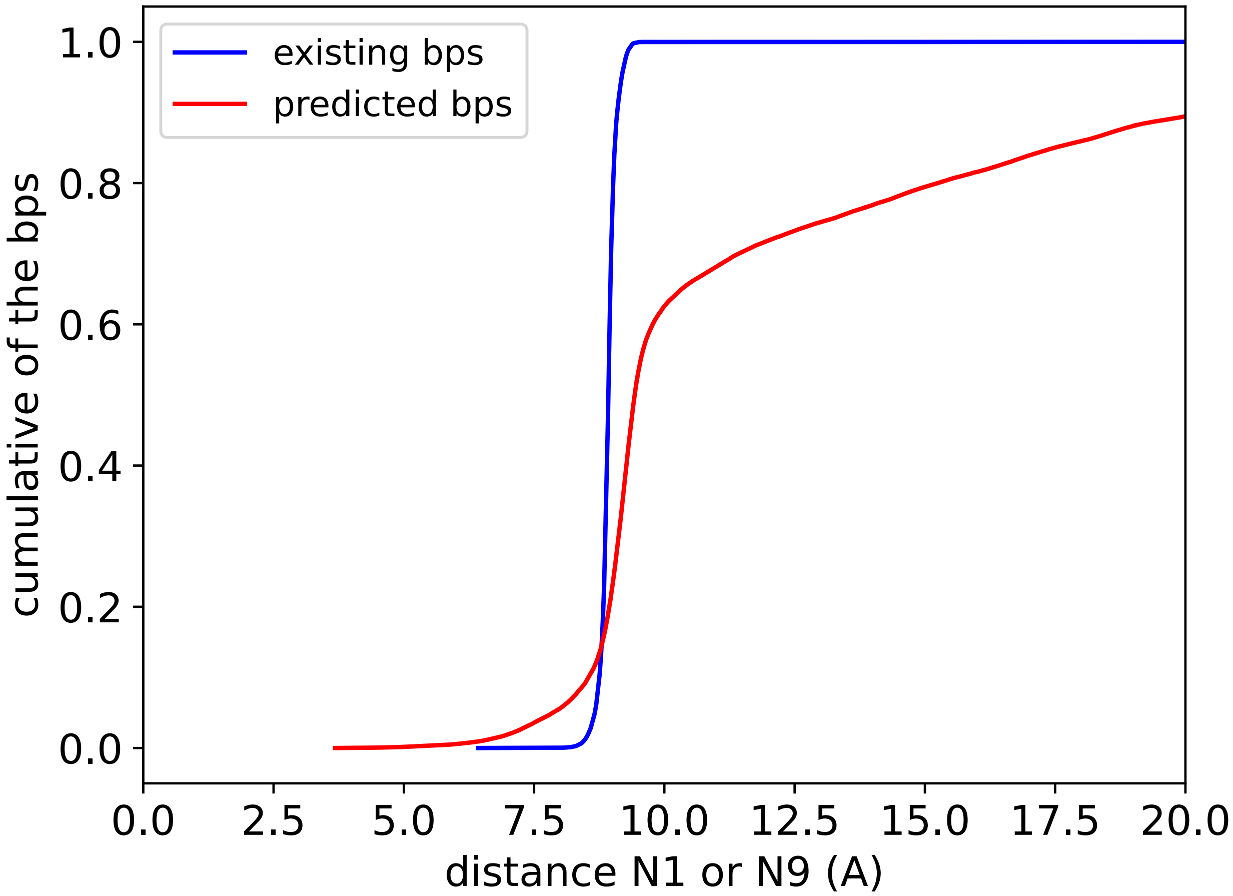


Figure S10: Cumulative distributions of distances between N1 or N9 among bases that are predicted to be paired: in blue the ones that were already present in the PDBs, in red the ones that were fully predicted. Around 80% of the pairs have a distance <15 Å, which is supposed to be a good distance to form a base pairing.
